# Hiding in the bush, in fear of a predator! Vegetation and predators influence shoaling among wild zebrafish

**DOI:** 10.1101/2023.07.02.547402

**Authors:** Ishani Mukherjee, Anuradha Bhat

## Abstract

Social responses of aquatic group-living organisms are critically dependent on predation risk and vegetation cover in their habitat. To gain insight into how these factors shape their immediate responses, we exposed wild zebrafish shoals to either vegetation, predator cues or both these factors simultaneously. Control treatments were not exposed to the above factors. By analyzing 60 unique shoals across 270 experimental trials, we found that while shoals formed significantly larger subgroups and were significantly more polarized in the presence of predator cues, both these properties decreased when shoals experienced predator cues in presence of vegetation. Furthermore, foraging was significantly lower when predator cues and/or vegetation were present. Tracking of all individuals in treatments devoid of vegetation revealed that: (i) compared to control treatments, individuals within shoals receiving predator cues had a significantly higher probability to continue being in a group and (ii) individuals occupying the front positions deviated lesser from their median position within a shoal as compared to other individuals. Anti-predator responses and foraging are critical for survival and therefore, this study provides important insights into shoal dynamics in changing environments.

## Introduction

Across taxa, ecological factors such as predation, resource availability and habitat complexity shape behaviour [1-5]. Animals show changes in behaviour as a response to immediate as well as long-term changes in their environment [6-7]. In the current study, we examine immediate shoal dynamic responses of wild-caught zebrafish towards two ecological factors: predation and vegetation cover. Findings from the current study not only increases our understanding on shoal plasticity towards different ecological cues but also can be applied in conservation strategies that involve regulating behaviour through altering ecological factors.

In shoaling fishes, antipredator responses and their ability to forage efficiently are directly linked to their survival [8-10], and hence, shoals are likely to exhibit considerable plasticity in behaviour in the presence of a predator or during foraging. Here, to gain insight into plasticity in immediate responses exhibited by shoals, we recorded the responses of wild zebrafish shoals (from the Ganges drainage in West Bengal, India) when exposed to predation threat under different ecological contexts. While populations of this cyprinid species are known to differ in behaviour based on predatory pressure, vegetation, and flow rate [11-13], behavioural plasticity exhibited in terms of shoaling and foraging by a single population is unclear.

We hypothesized that in presence of predator cues, individuals would disperse significantly lesser from group members. While shoals would adhere to safety in numbers in open habitats, in the presence of vegetation, individuals would take refuge underneath vegetation-a common antipredator response in fishes [14-15]. We further hypothesized that foraging would reduce in presence of predator cues as a) individuals would engage in antipredator behavior and b) in presence of vegetation, due reduced visual information on the presence of food. Fish schools are often led by certain individuals (known as leaders) and we expected similar behavior in zebrafish shoals.

## Methods

### Collection and maintenance of fish

Wild zebrafish shoals were collected from the Ganges drainage basin in West Bengal in December 2019 and January 2020 (Habitat specifics: Supplementary Material, S1.1). Shoals were brought to the laboratory and were maintained in bare, aerated 60litre tanks (100-120 individuals/ tank). *Channa* spp. (snakeheads) individuals were also collected from the same habitat (Mean length:12cm), brought to the laboratory and were kept in 18litre tanks (1 individual/ tank). A temperature range of 23°C-25°C and a constant lighting condition of 12h dark:12h light in the laboratory was maintained. While the zebrafish were fed daily ad-libitum with freeze-dried bloodworms or brine shrimp (*Artemia spp*.), the snakeheads were fed daily with pellet food or zebrafish that died of natural causes.

### Experiments

Shoal responses were recorded under four treatments: in presence of predator cues, vegetation or both ecological factors. In control treatments (C), shoals were recorded devoid of predator cues and vegetation. In predator cue treatments (or PT), shoals were recorded in the presence of olfactory cues from their natural predator, the snakehead (*Channa* spp.). To simulate the presence of a predator, 6.5l of water with predator olfactory cues was added gently into the center of the arena [16-20] (Supplementary Material, S1.2). In vegetation treatments (or VT), shoals received visual stimuli of vegetation in the form of three identical plastic aquarium plants placed on diagonally opposite corners of the arena. In treatments subjected to both predator cues and vegetation (or PVT), shoals were introduced into an arena with vegetation and were provided olfactory cues from a predator as in PT.

Experiments were performed between 11:00-15:00 hours. In their natural habitats, comprising shallow ditches, wild zebrafish typically form shoals comprising 10-20 individuals [21] and thus, a shoal comprising ten individuals was gently transferred into a 75cm×75cm×12cm arena filled with aged water. A water depth of 5cm was maintained across treatments. Shoals exposed to olfactory cues from the predator were put in the arena with water depth ∼4cm as the final water depth would become 5cm after addition of olfactory cues. To avoid the impact of sex of individuals on shoaling behaviour [22-23], we randomly chose individuals who constituted a shoal and thereby maintained the population sex ratio of roughly equal number of males and females. Ground vibrations were minimized by placing a 3.5cm thermocol sheet layer beneath the arena. Two 20W LED light bulbs on either side of the arena maintained a constant light source. Prior to recording, the shoal was allowed to acclimatize in the arena for 20 minutes. Two minutes after addition of cues, (to allow shoals to recover from the disturbance caused by addition of cues - if any), or immediately after acclimatization in case of VT or C, the shoal was video recorded for 20 minutes at a resolution of 25 frames per second, using a camera (Canon Legria HF R306) placed overhead. Following this, 0.25g of freeze-dried blood worms were introduced into the arena center and the shoal was video recorded for another 5 minutes. The arena was emptied and rinsed with aged water between consecutive trials to remove cues from blood worms, conspecifics or of a predator. A total of 60 unique shoals were tested – 30 shoals were tested in all four conditions and 30 more shoals were tested either in PT, and C conditions (15 shoals/ condition). Trials were performed in a random order and all analyses of video recordings was performed by a single observer (IM) blind to the treatment.

### Statistical Analysis

Shoaling behaviour was quantified by measuring the size of the largest subgroup and the polarization state of shoals every 30s (750 frames) for 20 minutes. Individuals were considered to be within a subgroup when any part of their body was within two body lengths from one another, and the size of the largest subgroup was noted. The polarization score at a given frame was measured as the fraction of the number of individuals aligned in the same direction per number of individuals in the shoal [24-25]. Therefore, while a polarization score of 0 indicates no alignment among fishes, a polarization score of 1 indicates that all fishes are aligned in the same direction. To check whether these parameters were consistent over time, the 20-minute videos were divided into four 5-minute sessions (termed as Session 1, Session 2, Session 3 and Session 4). Thereafter, the mean largest subgroup size and the mean polarization score for each session was calculated. To estimate foraging across the four treatments we manually counted the number of bites at bloodworms by each shoal in the first two minutes of the recording.

All individuals belonging to C and PT treatments were tracked for the first five minutes using idTracker and thereafter errors in their trajectories (if any) were manually corrected to reach a tracking accuracy of almost 100% [26]. A total of 30 shoals (15/treatment) were tracked. Tracking was done in treatments where vegetation was absent as knowing the precise position of fish underneath vegetation was not feasible. Following Borner et al. [27], and Krause and Ruxton [28] we assigned solitary or group states to individuals every 10s (250 frames) (Section S1.3, Supplementary Material). The probability of not switching states i.e., remaining solitary and remaining in group state was calculated. Next, following Doughty et al. [29], we manually noted down the movement order of individuals within the largest subgroup every 10s (250 frames). The leader of the largest subgroup (of size *n*) received place 1, the next closest received position 2, and so forth. The last position was *n*. Following Flischoff et al., 2017 [41], we normalized for variations in largest subgroup size by calculating their position index using the following formula:

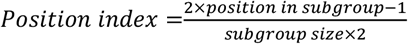

The median position index of individuals was thereafter calculated (Section S1.4, Supplementary Material) and the standard deviation from their median position indexes were calculated using the following formula:

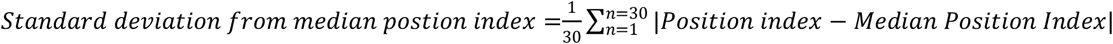

All analyses were performed using R Studio [30]. Generalized linear mixed models (GLMMs) were built (using ‘lme4’ and ‘lmerTest’) to understand the effect of: (i) treatment (CT, PT, VT or PVT) and session (Sessions 1-4) on largest subgroup size and polarization, (ii) treatment on percentage shoal under vegetation. In these GLMMs, shoal identity was incorporated as the random factor. Similarly, separate generalized linear models (GLMs) were built to understand the effect of treatment on: (i) foraging behaviour (number of bites at worms) and (ii) the probability of not switching states. Our data was not near to any distribution (checked using ‘fitdistr’ [31]), therefore we ran models assuming our data was normal. Model comparisons were performed using ANOVA in ‘car’ package [32] and post hoc paired tests (Tukey’s post hoc HSD Test using the ‘multcomp’ package) were performed for comparing the effects of factors that were significant. We performed Spearman’s Corelation to check the correlation between their median position index and median deviation. Mean ± standard error values have been reported throughout the manuscript. Two tailed p-values less than or equal to 0.05 were considered significantly different.

## Results

### Mean largest subgroup size, mean polarization score and foraging behaviour

The GLMM revealed that mean largest subgroup size was significantly impacted by treatment (Wald type IIχ^2^ = 163.65, df = 3, p<0.001) and that mean largest subgroup size was comparable across sessions (Wald type IIχ^2^ =1.79, df = 4, p=0.77; Table S1A). The mean largest subgroup size of PT shoals (5.72±0.13) was significantly greater than the mean largest subgroup size of C (4.06±0.09), VT (4.18±0.10) and PVT (4.64±0.12) shoals (Figure 1A). The GLMM revealed that mean polarization score was significantly impacted by treatment (Wald type IIχ^2^ = 187.82, df = 3, p<0.001) and that mean polarization score was comparable across sessions (Wald type IIχ^2^ =3.54, df = 3, p=0.31; Table S1B). The mean polarization score of PT shoals (0.56±0.01) was significantly greater than the mean polarization score of PVT (0.46±0.01), VT (0.38±0.01) and C shoals (0.41±0.01) (Figure 1B). The GLM revealed a significant effect of treatment (Wald type IIχ^2^ = 22.78, df = 3, p<0.001) on number of bites at worms (Table S1C): C shoals bit significantly more worms (30.93±2.31 bites) than shoals in VT (17.41±2.32 bites), PT (20.33±2.40 bites) or PVT (17.75±2.01 bites) (Figure 1C) (Tukey’s test results in Table S1D). The GLMM revealed a significant effect of treatment on percentage shoal under vegetation (Wald type IIχ2 = 3.96, df = 1, p=0.04) - a significantly greater percentage of individuals were under vegetation in PVT (18.23±1.58%) as compared to VT. (24.31±1.89) (Figure 1D, Table 1SC)

**Figure 1:**
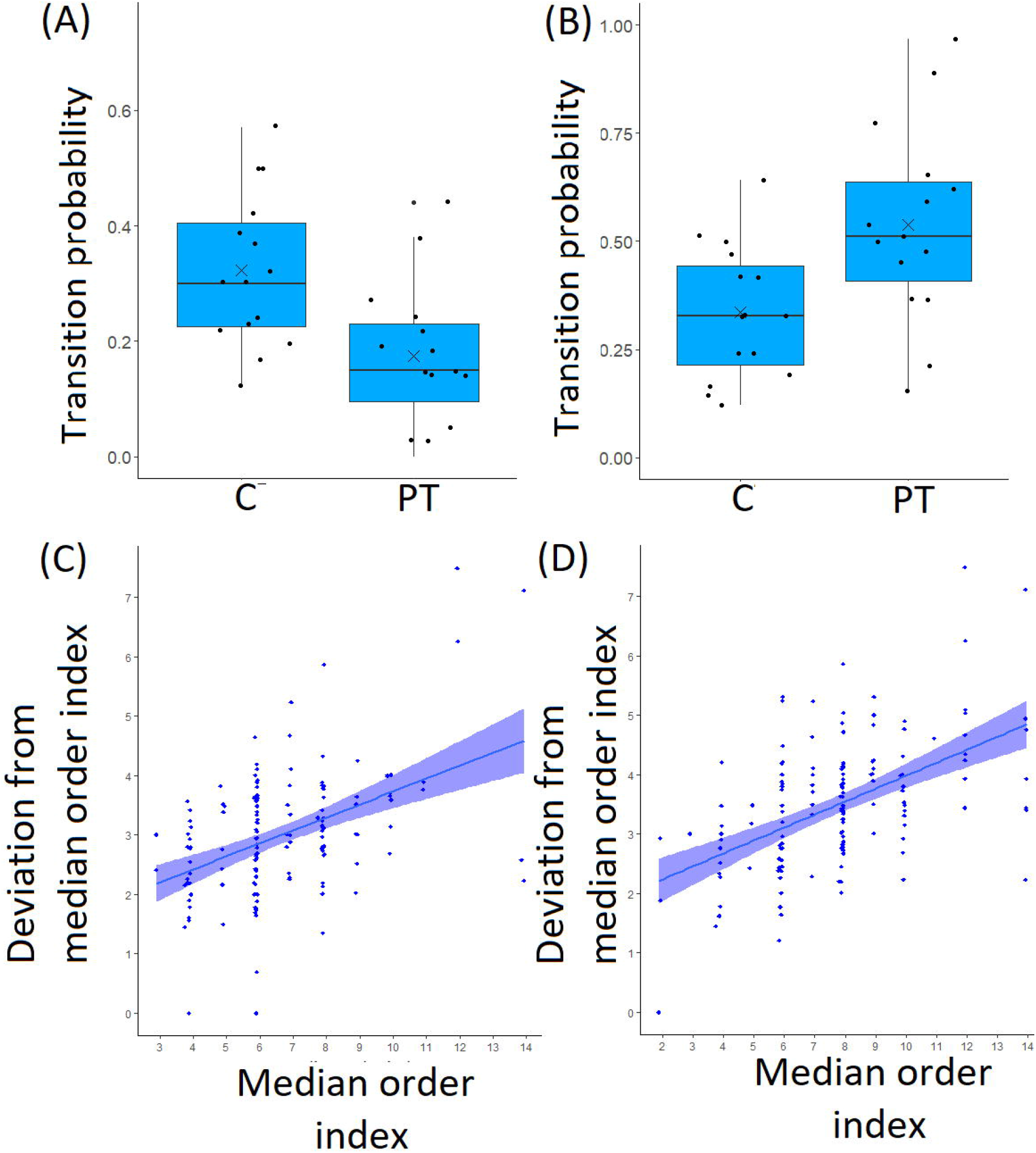
Box-and-whisker plots across treatments representing: (A) The mean size of largest subgroup, (B) The mean polarization score and (C) Number of bites at worms. Each data point is represented as a dot. The different letters placed above the boxes represent significant differences between the categories. Comparisons were performed using Tukey’s HSD Test (Sample size-Mean size of largest subgroup and mean polarization score: N_C_ = N_PT_ = 45 shoals; Nv_T_ = N_PVT_ =30 shoals; p<0.05)

### Shoal dynamics within shoals receiving cues from a predator

The probability of continuing to swim solitary or continuing to swim in a group was dependent on the treatment (GLM results for: (1) continuing to swim solitary: Wald type IIχ^2^ = 10.03, df = 1, p>0.01, Table S2A; (2) continuing to swim in group: Wald type IIχ^2^ = 8.13, df = 1, p>0.01; Table S2B). The probability of continuing to be in solitary state was significantly smaller among individuals in PT shoals (0.17 ±0.03) as compared to individuals in CT shoals (0.32±0.01) (Figure 2A). Correspondingly, the probability of continuing to be in a group was significantly higher among individuals in PT shoals (0.53±0.05) as compared to individuals in CT shoals (0.33±0.03) (Figure 2B) (Tukey’s test results in Table S2C).

**Figure 2:**
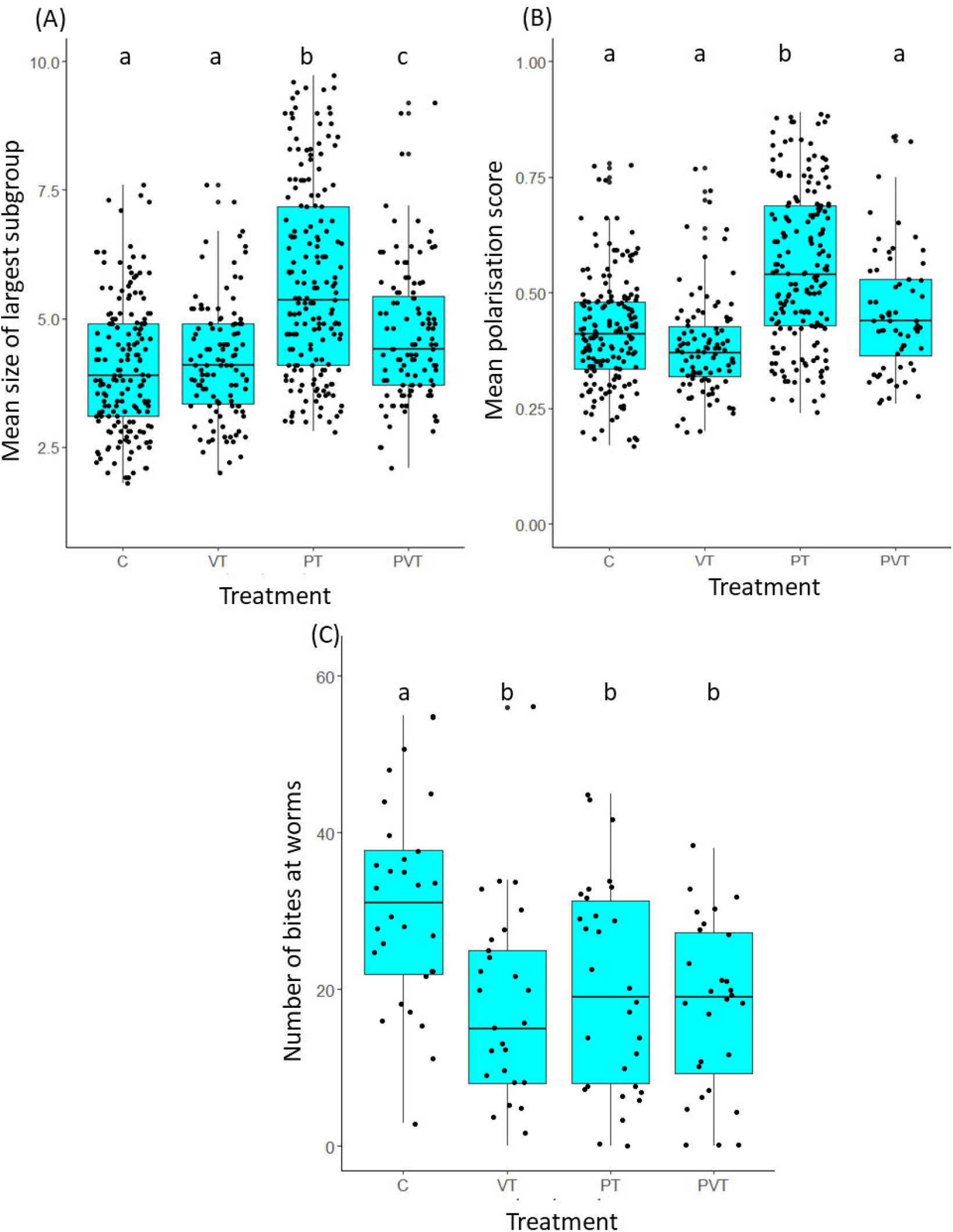
Shoal dynamics and deviation in individuals’ shoal position in control treatments and treatments receiving predator cues. Box-and-whisker plots representing the probability of continuing to swim in: **(A)** solitary and **(B)** in group state. Each data point is represented as a dot. The different letters placed above the boxes represent significant differences between the categories. Comparisons were performed using Tukey’s HSD Test. (Sample size-N_C_ = N_PT_ = 15 shoals; p<0.05). Individual variation of fish position: median position of each fish plotted against the standard deviation from the median position for **(C)** shoals in control treatments and **(D)** shoals predator cue treatments. (Sample size-N_C_ =140 individuals; N_PT_ = 130 individuals)

There was a strong correlation between median position and the standard deviation from the median position for both treatments (C: R^2^=0.43; p<0.001; PT: R^2^=0.53; p<0.001). Individuals in PT shoals and individuals in C shoals showed a similar pattern with regards to their shoal position: individuals towards the front deviated lesser from their median position in the shoal as compared to other individuals (Figure 2C and Figure 2D).

## Discussion

We found that wild zebrafish shoals show considerable plasticity in shoaling and foraging response in the presence of predation and vegetation. Consistent with studies on mosquitofish (*Gambusia affinis*), guppies (*Poecilia reticulata*), wild piranha (*Pygocentrus nattereri*) and fathead minnows (*Pimephales promelas*) [34-37], shoal cohesion increased as individuals came close to one another to form larger subgroups in presence predator odour cues, and more cohesive groups also exhibited increased polarization [38-39]. As observed across taxa [40-42], the presence of vegetation reduced subgroup size and polarization as shoals switched evasion strategy from safety in numbers to seeking refuge to escape threat.

Ecological factors like habitat, prey availability and predation strongly control fission-fusion dynamics among schooling fishes [43-45]. As in guppies, the analyses of fission-fusion dynamics revealed that in the presence of predator cues, the tendency of individuals to leave the largest subgroup declined significantly [46]. In fish species such as the Atlantic cod (*Gadus morhua*), gold shiners (*Notemigonus crysoleucas*), mosquitofish (*Gambusia affinis*), guppies (*Poecilia reticulata*) specific individuals (termed as leaders) consistently occupy the front of a shoal. Our results also reveal that individuals towards the front of a zebrafish shoal showed lesser deviation from their positions as compared to individuals who followed, suggesting that shoal leaders consistently lead the shoal over the observation period.

While shoals forage most effectively in the absence of vegetation and predation, a reduction in foraging in presence of a predator, possibly as an antipredator response [48-50] in presence of vegetation, could be explained by the following hypotheses. Owing to the fact that zebrafish are visual foragers [51-52], the presence of vegetation is likely to disrupt the access to visual information about the presence/location of food. Therefore, this lack of/ reduced information might cause reduced foraging in an environment with vegetation. Another possibility could be that test shoals might choose a refuge (in the form of vegetation) over foraging in the open arena. Increasing the shoals’ motivation to forage by starving fishes for 24 hours prior to foraging trials will enable us to decipher whether it is the latter factor.

In their natural habitats, zebrafish often feed on vegetation, algae and zooplanktons found in the waterbodies [53] and therefore in such habitats, vegetation and their food sources are often not spatially separated. While this study clearly shows that vegetation may be useful in the context of predator avoidance, the same factor may obstruct useful information such as the presence of food sources. Thus, this study may provide useful insights to conservation efforts involving provision of vegetation cover along banks of stream/river/lakes.

## Supporting information

Electronic Supplementary File

## Data accessibility

Data for this study are available at https://datadryad.org/stash/share/DFpf3Ab-7Kyub9JdmVAGn-9dg0wveTwfO0oW9xNaODo

## Author contributions

I.M. and A.B. conceived the study and designed the experiments. I.M. carried out all the experiments and the data analysis. I.M. and A. B. wrote the manuscript.

## Acknowledgements

The authors thank the Indian Institute of Science Education and Research Kolkata (IISER Kolkata), India, for providing infrastructural and financial support. The authors also thank Prasenjit Pan for help in collection of the wild population and fish maintenance in the laboratory.

## Ethics Statement

Guidelines outlined by the Committee for the Purpose of Control and Supervision of Experiments on Animals (CPCSEA), Ministry of Fisheries, Animal Husbandry and Dairying, Government of India were followed in all aspects of maintenance and experimentation. All experimental protocols followed here have been approved by the Institutional Animal Ethics Committee’s (IAEC) and guidelines of Indian Institute of Science Education and Research (IISER) Kolkata, Government of India (Approval number IISERK/IAEC/AP/2021/70).

