## Supplementary material for "Hiding in the bush, in fear of a predator! Vegetation and predators influence shoaling among wild zebrafish": Electronic Supplementary File

**S1.1 Habitat specifics**

| **Parameter** | **Range/ Description** |
|---|---|
| Temperature | 23°C-30°C |
| pH | 6.8-8.4 |
| conductivity | 460µS/cm - 500 µS/cm |
| TDS | 210ppm-265ppm |
| Vegetation | Moderate vegetation consisting of submerged grasses, and shrub plants was prevalent along the edges of the habitat |
| Most abundant predator | *Channa spp.* (snakeheads), a common predator to zebrafish, also occurred in the same habitat. |

**S1.2 Detailed protocol for addition of predator cue in PT and PVT**

Prey species elicit anti-predator responses based on previous exposure to predators [16-18]. In the current study, test shoals were wild-caught and thus, prior to their collection, these shoals have experienced odour cues from predators (commonly, *Channa* spp.). Therefore, zebrafish shoals used in our study would recognize the danger of predator cues and would elicit anti-predator responses. Previous studies show that water from a tank housing a predator contains olfactory cues from the predator that evokes antipredator responses in prey [19-20]. Based on the above facts, water from a *Channa* tank was added in treatments simulating the presence of a predator. Control experiments established that gentle addition of water into the arena center had no impact on shoaling properties (unpublished data)

**S1.3 Details for categorizing individuals into solitary or group state**

Following Borner et al., we recorded the position of each fish and that of its neighbours every 10s (250 frames) [27]. Individuals who had other individual(s) within four body lengths were considered to be in a group [27, 28] and individuals who did not have so were considered to be solitary. The rationale for setting the criteria of being within four body lengths to be in group state and setting the criteria of being within two body lengths to be a part of a subgroup is as follows: four body lengths is a looser cut off and is optimum (as supported by other studies on schooling fish) to categorize whether an individual is within a group or is alone. On the other hand, two body lengths is a more stringent cut off, and therefore, was has been used to assign individuals into subgroups.

**Table S1A:** Results of the generalized linear mixed model (GLMM) for predicting effect of treatment and session on mean largest subgroup size

Model: Mean size of largest subgroup ~ session + treatment + (1|shoalid)

Coefficients:

| | Estimate | Std. Error | t value | Pr(>\|t\|) |
|---|---|---|---|---|
| (Intercept) | <0.001 | <0.001 | 13.36 | <0.0001 |
| Session 1 | <0.001 | <0.001 | -0.01 | 0.99 |
| Session 2 | <0.001 | <0.001 | 0.64 | 0.52 |
| Session 3 | <0.001 | <0.001 | 0.30 | 0.76 |
| Session 4 | <0.001 | <0.001 | 0.19 | 0.85 |
| PT | <0.001 | <0.001 | 11.23 | <0.001 |
| PVT | <0.001 | <0.001 | 3.20 | <0.01 |
| VT | <0.001 | 0.001 | -0.01 | 0.99 |

**Table S1B:** Results of the generalized linear mixed model (GLMM) for predicting effect of treatment and session on mean polarization score

Model: Mean polarization score ~ session + treatment + (1|shoalid)

Coefficients:

| | Estimate | Std. Error | t value | Pr(>\|t\|) |
|---|---|---|---|---|
| (Intercept) | 0.41 | 0.01 | 23.24 | <0.0001 |
| Session 1 | 0.02 | 0.01 | 1.82 | 0.98 |
| Session 2 | 0.01 | 0.01 | 0.93 | 0.62 |
| Session 3 | 0.01 | 0.01 | 0.55 | 0.79 |
| PT | 0.14 | 0.01 | 11.49 | 0.86 |
| PVT | 0.02 | 0.01 | 1.40 | <0.0001 |
| VT | -0.01 | 0.01 | -1.18 | <0.0001 |

**Table S1C:** Results of the generalized linear model (GLM) for predicting effect of treatment on number of bites at prey (blood worms)

Model: Number of bites ~ treatment

Coefficients:

| | Estimate | Std. Error | t value | Pr(>\|t\|) |
|---|---|---|---|---|
| (Intercept) | 30.93 | 2.28 | 13.51 | <0.0001 |
| PT | -10.60 | 3.23 | -3.27 | 0.001 |
| PVT | -13.18 | 3.29 | -4.00 | <0.001 |
| VT | -13.52 | 3.26 | -4.14 | <0.0001 |

**Table S1D:** Tukey's test results for mean size of largest subgroup, mean polarization score and number of bites at blood at prey (blood worms)

*Mean size of largest subgroup*

| | Estimate | Std. Error | Z value | Pr(>\|z\|) |
|---|---|---|---|---|
| PT - C | 1.55 | 0.13 | 11.24 | <0.001 |
| PVT - C | 0.46 | 0.14 | 3.21 | <0.01 |
| VT - C | -0.01 | 0.15 | -0.003 | 1.00 |
| PVT - PT | -1.09 | 0.14 | -7.76 | <0.001 |
| VT - PT | -1.55 | 0.15 | -10.09 | <0.001 |
| VT - PV | -0.46 | 0.15 | -2.89 | 0.02 |

*Mean polarization score*

| | Estimate | Std. Error | Z value | Pr(>\|z\|) |
|---|---|---|---|---|
| PT - C | 0.14 | 0.01 | 11.49 | <0.001 |
| PVT - C | 0.02 | 0.01 | 1.40 | 0.49 |
| VT - C | -0.01 | 0.01 | -1.18 | 0.63 |
| PVT - PT | -0.12 | 0.01 | -7.62 | <0.001 |
| VT - PT | -0.16 | 0.01 | -11.67 | <0.001 |
| VT - PVT | -0.03 | 0.01 | -2.38 | 0.07 |

*Number of bites at prey*

| | Estimate | Std. Error | Z value | Pr(>\|z\|) |
|---|---|---|---|---|
| PT - C | -10.60 | 3.23 | -3.27 | <0.01 |
| PVT - C | -13.18 | 3.29 | -4.00 | < 0.001 |
| VT - C | -13.51 | 3.26 | -4.14 | < 0.001 |
| PVT - PT | -2.58 | 3.29 | -0.78 | 0.86 |
| VT - PT | -2.91 | 3.26 | -0.89 | 0.80 |

| | | | | |
|---|---|---|---|---|
| VT - PV | -0.33 | 3.32 | -0.10 | 0.99 |

**Table 2A:** Results of the generalized linear model (GLM) for predicting effect of treatment on mean transition probability for continuing to be in solitary state:

Model: Mean transition probability ~ + treatment

| | Estimate | Std. Error | t value | Pr(>\|t\|) |
|---|---|---|---|---|
| (Intercept) | 0.32 | 0.03 | 9.70 | <0.0001 |
| PT | -0.14 | 0.04 | -3.16 | 0.01 |

**Table 2B:** Results of the generalized linear model (GLM) for predicting effect of treatment on mean transition probability for continuing to be in group state

Model: Mean transition probability ~ + treatment

| | Estimate | Std. Error | t value | Pr(>\|t\|) |
|---|---|---|---|---|
| (Intercept) | 0.33 | 0.04 | 6.72 | <0.0001 |
| PT | 0.20 | 0.07 | 2.85 | 0.008 |

**Table 2C:** Table showing the standard deviation of individuals from the frontmost, rearmost and median positions within a shoal.

| Treatment | Mean of position | Corresponding standard deviation |
|---|---|---|
| C | frontmost (3.83) | 2.57 |
| C | median position (7.48) | 3.83 |
| C | rear most (10.70) | 1.59 |
| PT | frontmost (3.35) | 2.77 |
| PT | median position (6.32) | 4.61 |

| PT | rear most | (11.00) | 2.36 |
|---|---|---|---|

**Table S1D:** Tukey's test results for continuing to be in a given state:

*Continuing to be in solitary state*

| | Estimate | Std. Error | Z value | Pr(>\|z\|) |
|---|---|---|---|---|
| PT - C | 0.14 | 0.04 | -3.17 | <0.01 |

*Continuing to be in group state*

| | Estimate | Std. Error | Z value | Pr(>\|z\|) |
|---|---|---|---|---|
| PT - C | 0.20 | 0.07 | 2.85 | <0.01 |
